# Inositol polyphosphates regulate resilient mechanisms in the green alga *Chlamydomonas reinhardtii* to adapt to extreme nutrient conditions

**DOI:** 10.1101/2024.09.17.613215

**Authors:** Rodrigo Bedera-García, María Elena García-Gómez, José María Personat, Inmaculada Couso

## Abstract

In the actual context of climate changing environments, photosynthetic organisms need to adapt to more extreme conditions. Microalgae can be excellent organisms to understand molecular mechanisms that activate survival strategies under stress. *Chlamydomonas reinhardtii* signaling mutants are extremely useful to decipher which strategies they use to cope with changeable environments. In this study, we conducted prolonged starvation in wild type and *vip1-1 Chlamydomonas* cells. The mutant *vip1-1* has an altered profile of pyroinositol polyphosphates (PP-InsPs) which are signaling molecules present in all eukaryotes. These molecules have been connected to P signaling in other organisms including plants but their implications in other nutrient signaling is still under evaluation. After prolonged starvation, WT and *vip1-1* showed important differences in the levels of chlorophyll and photosystem II (PSII) activity. We also performed a metabolomic analysis under these conditions and found an overall decrease in different organic compounds such as amino acids including arginine and its precursors and tryptophan which is considered as a signaling molecule itself in plants. In addition, we observed significant differences in RNA levels of genes related to nitrogen assimilation that are under the control of NIT2 transcription factor. Overall our data indicate an important role of PP-InsPs in the regulation of nutrient starvation especially regarding N assimilation and C distribution. These data are of great importance for the generation of resilient strains to be used in open ponds and high capacity bioreactors.

## Introduction

In the recent years, environmental challenges derived from uncontrolled CO_2_ emissions and other contamination sources have become a global priority. Improving water management and combating climate change have been included in the 2030 Agenda (https://sdgs.un.org/2030agenda) and SDGs of United Nations to transform our planet. Therefore, it is interesting to find sustainable solutions including the use of green organisms to address these problems. In this context, the use of microalgae stands as an interesting solution to achieve a reduction in greenhouse gas emissions through CO_2_ fixation in photobioreactors, and sustainable wastewater treatment (Kaloudas et al., 2021; Plöhn et al., 2021; Raeesossadati et al., 2014). Although we need to scale up and standardize the use of these microorganisms, we also need to understand the molecular mechanisms that make them adapt to environmental unfavorable conditions. In this way, *Chlamydomonas reinhardtii* is a single-celled green microalga that has been extensively used in different scientific contexts due to its well-characterized genetics and easy cultivation (Harris, 2009; Merchant et al., 2007). It has served as a model organism for nutrient-deprivation responses, such as carbon limitation (Wang et al., 2015), phosphorous deprivation (Couso et al., 2020; Hidayati et al., 2019; Kamalanathan et al., 2016; Zúñiga-Burgos et al., 2024), sulfur deprivation (De Mia et al., 2019; González-Ballester et al., 2010; Grechanik et al., 2020; Kajikawa et al., 2019) and nitrogen deprivation (Calatrava et al., 2019; Miller et al., 2010; Park et al., 2015; Saroussi et al., 2016; Schmollinger et al., 2014). Indeed, nitrogen assimilation mechanisms discovered in *C. reinhardtii* helped to elucidate how these processes are regulated in plants (Camargo et al., 2007; Castaings et al., 2009). To date, many efforts have been made to achieve biotechnological viability in using microalgae with waste products as a nutrient source (Behl et al., 2020; Han et al., 2023). However, in these applications, the nutrient input might fluctuate over time, leading to temporary limited nutrient conditions especially under prolonged cultivation. So that, it is important to improve our knowledge regarding nutrient limitation responses in microalgae to achieve optimal conditions in these biotechnological applications.

In *C. reinhardtii*, as well as in many other eukaryotic organisms, the Ser/Thr kinase Target of Rapamycin (TOR) is a conserved master regulator that integrates nutrient availability and energy levels to control central processes like amino acid, lipid and tetrapyrrole metabolism, nitrogen assimilation or carbon storage (González and Hall, 2017; Imamura et al., 2015; Kleessen et al., 2015; Mubeen et al., 2018; Pancha et al., 2020; Roustan and Weckwerth, 2018). In a previous study, the strain *vip1-1* was isolated in a rapamycin hypersensitivity screening of *C. reinhardtii* insertional mutants. This strain was deficient in highly phosphorylated inositol polyphosphate species, namely 1-diphosphoinositol 2,3,4,5,6-pentakisphosphate (1-PP-InsP_5_), 5-diphosphoinositol 1,2,3,4,6-pentakisphosphate (5-PP-InsP_5_) and 1,5-bis-diphosphoinositol 2,3,4,6-tetrakisphosphate (1,5-PP-InsP_4_) (Couso et al., 2016). These molecules are synthesized through consecutive phosphorylation of the *myo*-inositol and are usually referred to as inositol pyrophosphates (PP-InsPs), as they contain at least one pyrophosphate in their structure (Shears, 2015; Shears and Wang, 2020; Williams et al., 2015). The mutant strain *vip1-1* presents an insertion in the *vip1* gene which encodes an inositol hexakisphosphate kinase, responsible for the synthesis of inositol pyrophosphates in *C. reinhardtii* (Couso et al., 2016). PP-InsPs have been described as cell messengers which are involved in central signaling pathways mainly controlling protein phosphorylation, energy levels and P homeostasis in eukaryotic organisms including the green lineage (Mihiret et al., 2024; Nguyen Trung et al., 2022; Pipercevic et al., 2023; Saiardi, 2012; Wu et al., 2023). Interestingly, a synergistic signaling between TOR and PP-InsPs was described, uncovering a role of these molecules in carbon partitioning and tricarboxylic acid cycle control in *Chlamydomonas* (Couso et al., 2016). Moreover, a phosphoproteomic approximation using *vip1-1* strain revealed that the signaling network of the PP-InsPs exerts regulation over cellular energy levels and light stress response through regulation of non-photochemical quenching mechanisms in green algae (Couso et al., 2021). In the same work, it was also found that PP-InsPs affected the phosphorylation status of the photosynthetic apparatus, which leads to state transitions, further supporting the signaling role of these highly phosphorylated molecules in stress adaptations. Additionally, precursors of PP-InsPs have been shown to respond to CO_2_ levels in *Chlorella sorokiniana* and *Chlorella vulgaris* (Morales-Pineda et al., 2023) which is an important finding to further explore the biotechnological applications of this biosynthetic pathway.

Altogether, these reports highlight the emerging role of the inositol polyphosphates (InsPs) and especially the PP-InsPs in nutrient sensing and stress response. Therefore, in order to facilitate the applicability of microalgae in carbon sequestration and wastewater treatment, we decided to study the regulation of PP-InsPs over nutrient limitation responses in *C. reinhardtii*. To address this task, we employed a general nutrient deficiency by performing batch cultivation over an extended period of time, which allows us to capture different nutrient deprivation responses. Our results indicate that PP-InsPs regulate chlorophyll turnover and photosynthetic capacity under nutrient limitation. Moreover, we found that the metabolic profile of *Chlamydomonas* is deeply affected by the depletion of PP-InsPs in nutrient stressed cells, resulting in a remarked reduction in amino acids levels. Additionally, the transcriptomic data showed that these molecules are required for the nitrogen deprivation response controlled by the transcription factor NIT2. Therefore, this study provides new insights into the multi-layered regulatory effects of PP-InsPs regarding nutrient management and presents their biosynthetic pathway as potential genetic engineering target for biotechnological applications of green organisms.

## Materials and methods

### Strains and culture conditions

*Chlamydomonas reinhardtii* strain CC-1690 wild-type MT+ (Sager 21gr) (Sager, 1955) was used as the control strain, while the insertional mutant of the CC-1690 strain named *vip1-1* (Couso et al., 2016) was used as the object of study. The algal strains were grown at 25°C under 50 μmol m^-2^ s^-1^ photon flux density provided by OSRAM LED lamps. The experiments were initiated during exponential growth phase (1-2 x 10^6^ cells ml^-1^) in liquid TAP (tris acetate phosphate) medium, and performed after 24 hours (control), 14 or 21 days (nutritional stress) as stated for each experiment.

### Transmission electron microscopy

Cultures were collected at 1-2x10^6^ cells mL^-1^ and fixed with 2.5% glutaraldehyde in 0.1 M Na-cacodylate buffer at pH 7.4 for 2 h at room temperature. After fixation, cells were washed three times with the same buffer at room temperature. Samples were post-fixed in 1% osmium tetraoxide in cacodylate buffer (0.1 M, pH 7.4) for 1 h at 4°C. After washing, samples were immersed in 2% uranyl acetate, dehydrated through a gradient acetone series (50, 70, 90, and 100%), and embedded in Spurr resin (Spurr, 1969). Semi-thin 147 sections (300 nm thickness) were obtained with a glass knife and stained with 1% toluidine blue for cell localization and reorientation using a conventional optic microscope. Once a suitable block face of the selected area was trimmed, several ultrathin sections (70 nm) were obtained using an ultramicrotome (Leica UC7) equipped with a diamond knife (Diatome) and collected on 200 mesh copper grids. Sections were examined in a Zeiss Libra 120 transmission electron microscope and digitized (2048 × 2048 × 16 bits) using an on-axis mounted TRS camera.

### RNA-seq data generation and processing

Three independent biological replicates were considered for each strain and condition: wild-type strain after 24 hours (control) and 14 days of growth (nutritional stress), and *vip1-1* strain after 24 hours (control) and 14 days of growth (nutritional stress). RNA extraction was performed using mechanical disruption of the frozen cell pellets in a Mini Bead Beater (Biospe Products) mixed with 0.5 mm glass beads for individual cell lysis in the presence of an extraction buffer consisting of phenol:chloroform (1:1, v/v). RNA was purified using ISOLATE II RNA Plant Kit (Bioline) following manufacturer’s instructions. The quality of the RNA was firstly analyzed by gel electrophoresis and RNA integrity number (RIN) was computed using Agilent 2100 Bioanalyzer. RNA-seq data generation was obtained as described in Serrano-Pérez et al. (2022), and analyzed using the MARACAS tool (Romero-Losada et al., 2022). Approximately, 30 million 50 nt reads were obtained per sample. The RNA-Seq data was processed using fastQC (https://www.bioinformatics.babraham.ac.uk/projects/fastqc/) to perform the quality control, Hisat2 (Kim et al., 2019) to map reads, Stringtie (Pertea et al., 2015) to assemble the transcripts and Ballgown (Frazee et al., 2015) to quantify expression.

Raw data was normalized using the upper quantile method and low-accumulated genes (FPKM < 0.5 in all conditions) were pre-filtered. Differential expression analysis was performed using the Limma R package (Ritchie et al., 2015), based on linear models, and DESeq2 R package (Love et al., 2014), which relies on a negative binomial distribution. The minimum fold-change was set to 2 and the minimum adjusted p-value to 0.05 (BH method was used to adjust p-values). In histograms of gene expression, the values are displayed in Fragments Per Kilobase of transcript per Million mapped reads (FPKM) after quantile normalization.

### Metabolomic data generation and processing

Five independent biological replicates were assessed for each strain and condition: wild-type strain after 24 hours (control) and 14 days of growth (nutritional stress), and *vip1-1* strain after 24 hours (control) and 14 days of growth (nutritional stress). The metabolite content determination was performed using cell pellets that firstly were lyophilized (Skadi-Europe TFD 8503), and stored at −20°C as described in Serrano-Pérez et al. (2022). Metabolites were determined from 15-20 mg of lyophilized cell biomass subjected to mechanical disruption in a Mini Bead Beater (Biospe Products) with 0.5 mm glass beads in the presence of 1 mL extraction buffer consisting of chloroform:methanol (3:7, v/v). Centrifugation at 5000 × g for 5 min at RT was then performed and these two steps were repeated until the pellets were found colorless. Supernatants were concentrated using speed-vacuum (Eppendorf concentrator Plus) and then samples were resuspended using 200 µL of MiliQ water and submitted to UPLC/MS analysis as described in Mccloskey and Ubhi (2015). Raw data was normalized using 10 µM paracetamol as an internal standard and the dry weight of each sample. The significance was calculated using the Wilcoxon signed-rank test, using the “stats” R package (https://www.R-project.org/), and the p-values obtained were adjusted using the Benjamini-Hochberg method implemented in the “qvalue” R package (Storey et al., 2024). Individual metabolite graphics were obtained using ggplot2 R package (Wickham, 2016), while the heatmap was generated using MetaboAnalyst 6.0 web tool (Pang et al., 2024) after auto-scaling each metabolite. Hierarchical clustering of metabolites was performed using Ward’s method and Euclidean distance to group metabolites behaving similarly across different samples.

### Chlorophyll determination and activity assays

Two independent biological replicates and two technical replicates were used for chlorophyll determination and activity assays in all conditions tested. 1-2x10^6^ cells mL^-1^ cultures of wild-type and *vip1-1* were used as controls. Chlorophyll determination was perform collecting cells by centrifugation at 6500 x g, resuspended in methanol and incubated at 70°C for 15 minutes. The supernatant obtained was then centrifugated at 6500 x g and used to perform OD determination using Thermo Scientific Genesys 30 spectrophotometer. The concentration of chlorophyll in the different samples was then calculated by measuring the absorbance at 650 nm and 665 nm according to Mackinney (1941).

The enzymatic activity of nitrate reductase was calculated in the different samples according to Paneque et al. (1965) with slight modifications. Cells were permeabilized by shaking vigorously during 10 s with 20 µL of toluene per mL of culture. 0.2 mL of a Tris-HCl 0.5 M and KNO_3_ 50 mM solution was added to 0.2 mL of cell culture, together with 0.2 mL of FMN 1 mM and water to a 0.9 mL final volume. 0.1 mL of a solution containing 8 mg mL^-1^ of Na_2_S_2_O_4_ in NaHCO_3_ 95 mM was added to the preparation to start the reaction, and after shaking, the samples were incubated 10 minutes at 30°C. The reaction was stopped with 1 mL of a sulfanilamide preparation (10 g L^-1^ sulfanilamide, 200 mL HCl 37% v/v, and water to 1 L), 1 mL of N-(1-Naphthyl) ethylenediamine dihydrochloride at 200 mg L^-1^ and 2 mL of water. After 10 minutes at room temperature, absorbance is determined at 540 nm. The enzymatic activity of glutamine synthetase was calculated in the same conditions, according to Shapiro and Stadtman (1970). Activity is displayed in mU, which corresponds to the amount of enzyme necessary to catalyze one nanomole of substrate per minute.

### Chlorophyll fluorescence measurements

Batch cultures of WT and *vip1-1*cells were grown in control conditions and 21 days in TAP medium at 25°C. We used cells suspensions of 2x10^6^ cells ml^-1^ and the measurements were performed in triplicate. Chlorophyll *a* fluorescence was measured using a pulse-amplitude modulation fluorometer (DUAL-PAM-100, Walz; Effeltrich, Germany). The maximum quantum yield of PSII was assayed after incubation of the cell suspension in the dark for 20 min with constant stirring by calculating the ratio of the variable fluorescence, Fv, to maximal fluorescence, Fm, (Fv/Fm). Induction/recovery curve was performed by exposing dark-adapted samples to constant illumination at 50 μE m^-2^ s^-1^ light intensity for 7 min, followed by 6 min of darkness. Effective PSII quantum yield (Y(II)) was determined using saturating pulses of red light at 5000 μE m^-2^ s^-1^ intensity and 100 ms duration. Y(II) was then calculated by the DUAL-PAM-100 software according to the equations reported by Kramer et al. (2004). Error bars indicate standard deviation (SD) of the values obtained from experiments performed in triplicate.

## Results

### Inositol pyrophosphates affect chlorophyll accumulation under nutritional stress

The *Chlamydomonas reinhardtii* insertional mutant *vip1-1* was characterized by Couso and colleagues and described as a strain deficient in pyrophosphorylated inositol species (InsP_7_ and InsP_8_) (Couso et al., 2016). This mutant showed a very interesting phenotype when subjected to long limiting nutritional conditions under batch culturing acquiring a greener appearance than WT, despite displaying no differences in control conditions (Figure 1A). Accordingly, chlorophyll quantification revealed that the mutant strain had similar levels of these pigments compared to the wild-type in control conditions, but increased 45% their chlorophyll content when subjected to nutritional stress after 21 days of batch cultivation (Figure 1B). Chlorophylls interact with different light-harvesting complexes to allow light absorption and structural stabilization in the photosynthetic apparatus (Bujaldon et al., 2017; Wang and Grimm, 2021). Therefore, we proceeded to assess the quantum yield of the photosystem II of the wild-type and mutant strains under control conditions and nutritional stress. We also evaluated the degree of damage at PSII by performing induction/recovery curves in the same conditions and strains (Figure 1C and 1D). We found that PSII activity was moderately affected in the *vip1-1* strain under control conditions, suggesting PP-InsPs are important to maintain optimal photosynthetic activity. Additionally, under nutritional limiting conditions, the maximum quantum yield of PSII (Y(II)) from the *vip1-1* mutant strain was further reduced to 55% compared to wild-type (Figure 1C). Although Y(II) was affected in both strains after nutrient limitation, induction/recovery curves (Figure 1D) showed that WT showed a better recovery of PSII than *vip1-1*, suggesting an important downregulation of photosynthetic capacity of this mutant under these conditions. All these data indicate that PP-InsPs are regulating the coupling of chlorophylls degradation and photosynthetic performance of the algal cells under nutrient limitation that seems to have an important role on the adaptation to nutritional stress conditions for green microalgae.

**Figure 1:**
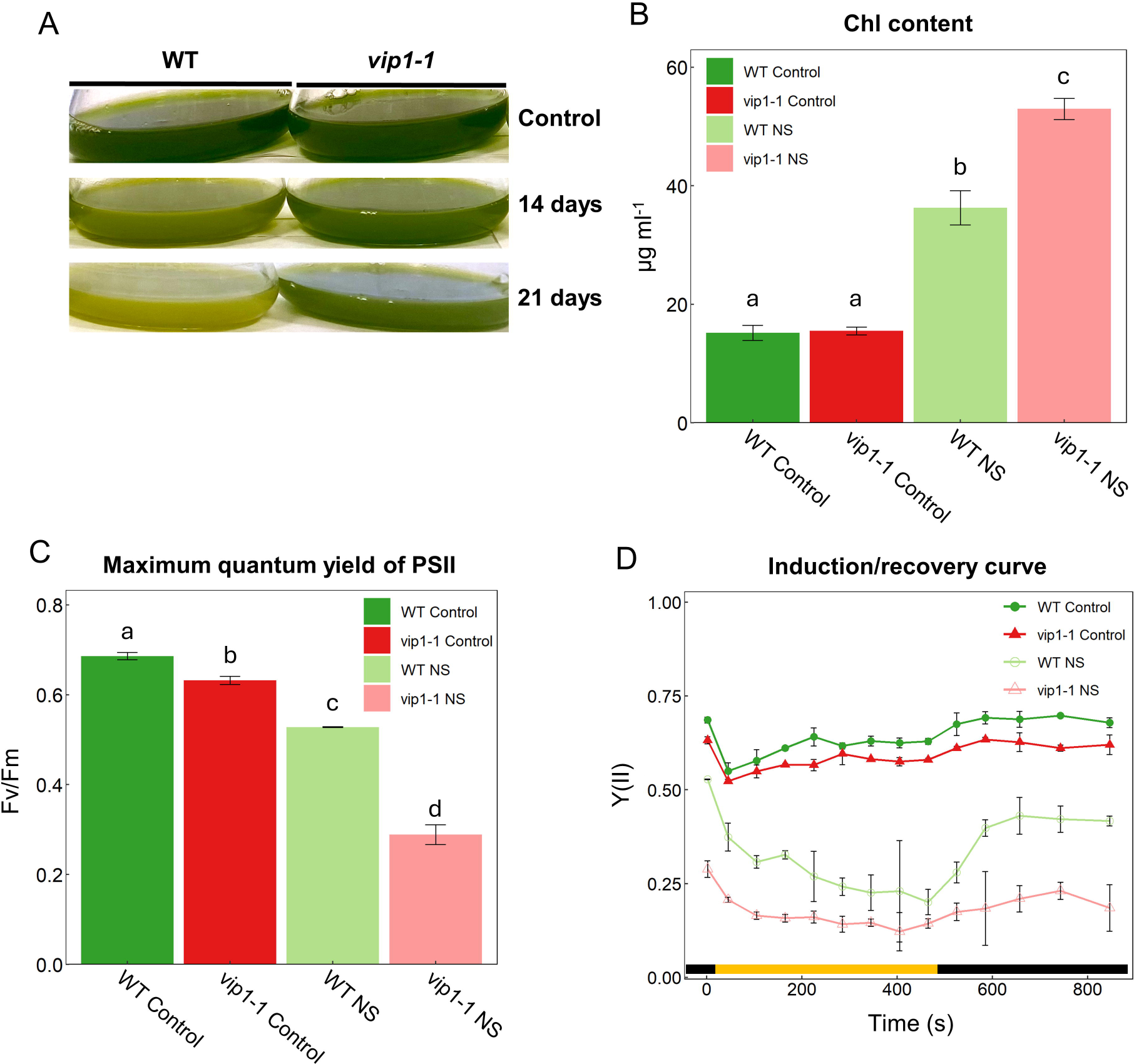
Appearance of cultures (A) after 24 hours (control), 14 and 21 days of batch cultivation (nutritional stress, NS). B and C, chlorophyll content (B) and maximum quantum yield of photosystem II (C) of WT and *vip1-1* strains after 24 hours (control) and 21 days of batch cultivation. D, Induction / recovery curve under control and NS conditions in WT and *vip1-1,* where the light exposure is represented with a yellow band. Significant differences are represented with different letters, evaluated using the Wilcoxon signed-rank test for B and Student’s t-test for C. Error bars indicate mean±SD.

### PP-InsPs influence ultrastructural adaptations to nutrient limitation

The increase in chlorophyll content together with limited Y(II) in a PP-InsPs deficient mutant under nutritional stress conditions (Figure 1B, 1C) may indicate that the *vip1-1* strain might show substantial differences at the ultrastructural level. We analyzed transmission electron microscopy images from wild-type and *vip1-1* strains after 1 day (Figure 2A, 2D), 14 days (Figure 2B, 2E) and 21 days (Figure 2C, 2F) of batch cultivation. These conditions mimic a complete nutritional limitation that not only depends on one major element, N or P for instance. After the TEM image analysis we found important structural differences in *vip1-1* cells compared to WT that became more evident as the nutritional limitation was longer. We observed two major differences between WT and *vip1-1* cells: the number of vacuoles and the electrodense material inside them. TEM sections containing *vip1-1* cells held between 10-12 vacuoles while we could only observe 3-4 vacuoles in WT cells after 1 day of culturing. After 14 and 21 days the number and electro density of the vacuoles in both strains were very similar especially after 21 days in batch. TEM analysis also showed some rounded structures inside vacuoles that could reflect a higher autophagic activity of *vip1-1* as described in Heredia-Martínez et al. (2018) in WT cells treated with cerulenin. These bodies are usually produced by the autophagic process under stress conditions, suggesting that PP-InsPs are involved in the regulation of autophagy together or in combination with TOR (Figure 2E) as we previously reported in Couso et al. (2021) when *vip1-1* were treated with TOR inhibitor, rapamycin. The second difference that we found after TEM analysis between WT and *vip1-1* cells was also the number and size of starch granules in *vip1-1* that started with very few starch granules and much smaller size (Figure 2, panels A and D) than WT under control conditions. However, after 21 days under batch cultivation, starch granules were very similar in number and size in both algal strains (Figure 2, panels C and F). These observations were confirmed by the analytical determination of starch (Supplementary figure 1) that shows an important increase in the starch levels in both algal strains after 21 days.

**Figure 2:**
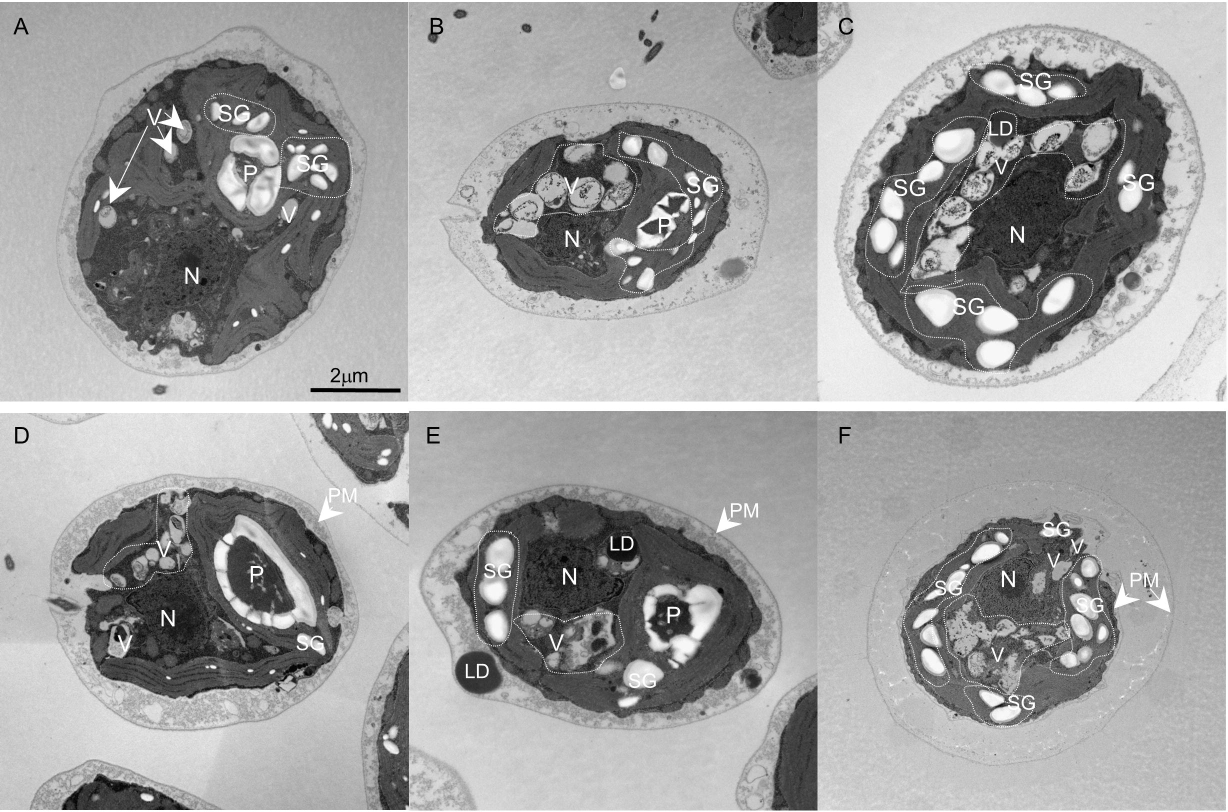
Electron microscopy **(TEM)** images from WT (A) and *vip1-1* **(D)** under control conditions. B and **E, TEM** images from WT and *vip1-1* after 14 days in batch cultivation respectively. C and **F, TEM** images from WT and *vip1-1* after 21 days in batch cultivation respectively. (V), vacuoles; **(P),** pyrenoid; **(N),** nucleus; (PM), plasma membrane; (SG), starch granules; **(LO),** lipid droplets.

Additionally, when we looked at the *C. reinhardtii* cells after 21 days of nutrient limitation, we found that *vip1-1* cells exhibited two plasmatic membranes and the wild-type cells did not, which suggests cell division control mechanisms might malfunction in nutrient-limited *vip1-1* cells (Figure 2C, 2F). In this context, previous work has proven that defects in autophagy reduces mating efficiency when subjected to nitrate deficiency (Kajikawa et al., 2019). The observed alterations in the autophagic process of *vip1-1* displayed by the TEM images after 14 days of nutritional limitation (Figure 2E), might be contribute to the problems detected in the division events in *vip1-1* cells after 21 days of batch cultivation (Figure 2F).

### Metabolomic analysis reveals effect of PP-InsP in N assimilation

In order to detect possible differences and deregulations in metabolic routes in *vip1-1* under nutritional limitation, we performed a metabolomic analysis of the wild-type and *vip1-1* strains under control and prolonged cultivation conditions. UPLC/MS was used (see methods section) to analyze over 50 metabolites belonging to different metabolic pathways. Changes in metabolites accumulation were captured using a heatmap plot (Figure 3A) generated by MetaboAnalyst 6.0 (Pang et al., 2024). A widespread decrease in metabolite content as a response to nutritional stress became evident for both the wild-type and the *vip1-1* strains. Remarkably, we noticed that the absence of PP-InsPs produced a strong effect over the amino acids content. In control conditions, *vip1-1* showed significantly higher levels of arginine, cystine, methionine, tyrosine, phenylalanine histidine, tryptophan and L-citrulline (Supplementary table I). When the cells were subjected to nutrient limitation, 12 and 2 amino acids were significantly downregulated and upregulated respectively in the *vip1-1* strain (Supplementary table I). This general increase in nutrient replete conditions and decrease in nutrient limitation suggests that the absence of PP-InsPs prevents the *vip1-1* strain from correctly adjusting to the metabolic demands. Tryptophan, arginine and L-citrulline stood out because they were higher accumulated in *vip1-1* under control conditions, but greatly reduced in nutrient stress conditions (approximately between 3- to 6-fold reduction) (Figure 3A; Supplementary table I). Although tryptophan functions as a regulatory compound to coordinate growth and stress responses in plants (Liu et al., 2022), little is known about its effect in microalgae besides translation. On the other hand, in *C. reinhardtii* the arginine biosynthetic pathway has been proposed to participate in responses to the nitrogen status. High arginine levels, through the production of nitric oxide and possibly other signaling pathways, lead to repression of nitrogen assimilatory mechanisms such as nitrate transporters and nitrate reductase (de Montaigu et al., 2010; González-Ballester et al., 2018; Monteiro et al., 2023). Interestingly, we detected at least a 2.5-fold reduction in the metabolites involved in the arginine biosynthesis including the intermediate L-citrulline in the *vip1-1* mutant compared to the wild-type strain under nutritional stress conditions (Supplementary table I, Figure 3B).

**Figure 3:**
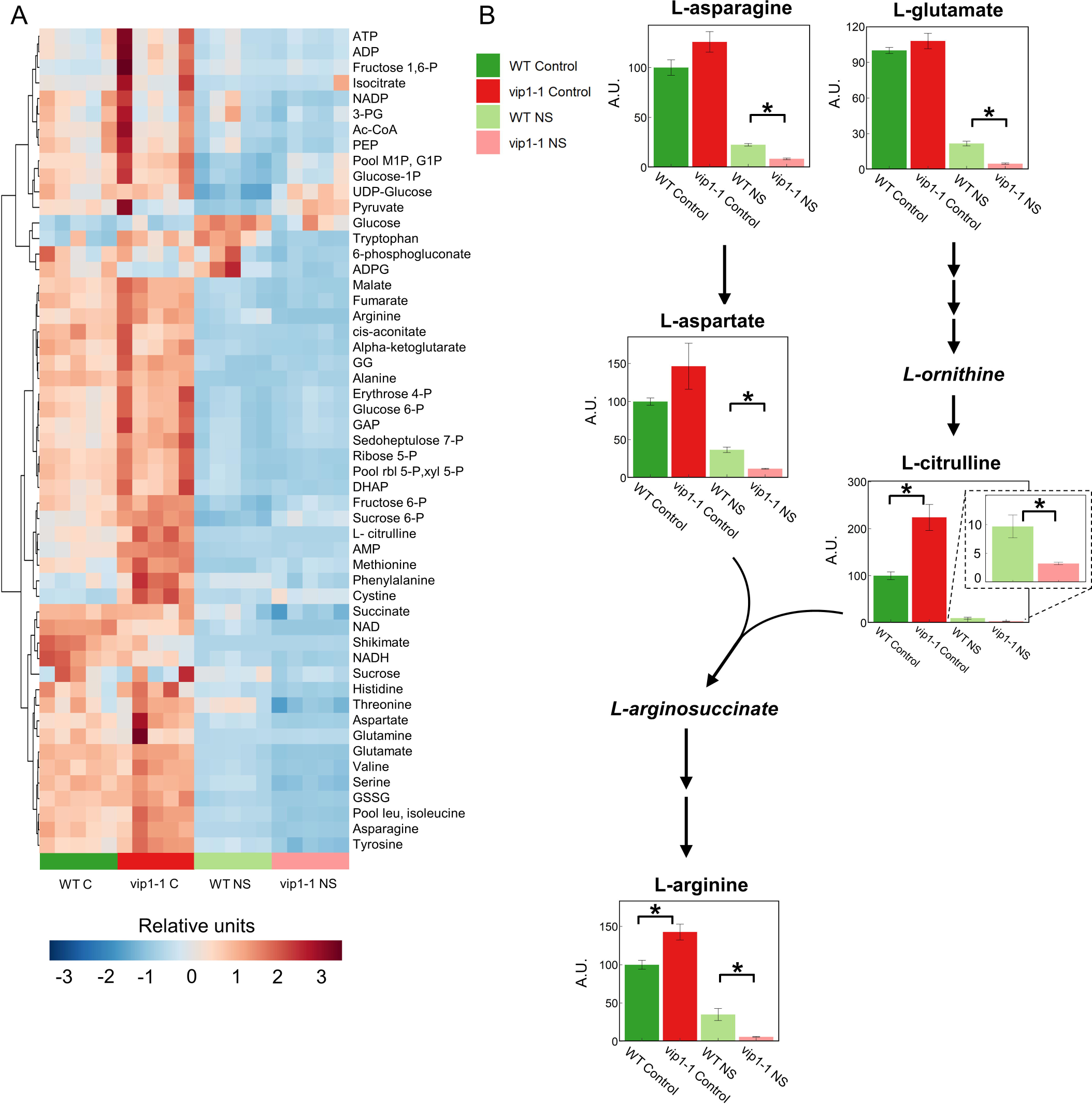
Heatmap of metabolites (A) representing their levels in each biological sample treated under control conditions (C) and nutrient starvation (NS). Row clustering was calculated using Ward’s mehtod and euclidean distance. B, diagram of the arginine biosynthetic pathway and the accumulation levels of some intermediates, relative to WT control levels. Consecutive arrows represent omitted intermediates, cursive letters indicate non-determined metabolites. Asterisks represent significant differences (adjusted p-value < 0.05), evaluated using Wilcoxon rank-sum test with the BH procedure adjustment. Error bars indicate mean±SE.

Additionally, pyruvate, glucose-1P and UDP-glucose grouped together using the clustering algorithm. They did not show differences under control conditions between strains but were significantly more abundant under nutritional stress in the *vip1-1* mutant strain (Supplementary table I). These compounds take part in central carbon biosynthetic pathways, like amino acid and tricarboxylic acid cycle intermediates (pyruvate), starch (glucose-1P) or glycerolipids (UDP-glucose) (Deschamps et al., 2023; Li-Beisson et al., 2023; Zhu et al., 2024). However, the glucose-1P must be transformed into ADP-glucose in order to synthesize starch out of glucose-1P, and we observed a significant 8-fold reduction in ADP-glucose accumulation in the *vip1-1* strain compared to the wild-type under nutritional deficiency (Supplementary table I). In these conditions, we also found an important decrease of starch levels in *vip1-1* compared to the wild-type in control conditions, but these differences are alleviated after 21 days (Supplementary figure 1).

These results suggest a role for inositol pyrophosphates in the sensing of nutrient availability, particularly nitrogen, together with carbon utilization possibly through the fine-tuning of the signaling processes that govern the adaptations of the central carbon metabolism. However, the mechanism by which the PP-InsPs tune these signaling processes is still unclear, as both upregulation and downregulation of metabolites are here identified.

### PP-InsPs participate in N deprivation response through a multilayered regulation

The metabolic profiling of the PP-InsPs deficient strain *vip1-1* under nutritional stress (NS) revealed a reduction in amino acids involved in the biosynthesis of arginine, and arginine itself. Previously, arginine levels have been proposed to control transcription levels of N assimilation-related genes (González-Ballester et al., 2018; Monteiro et al., 2023). Therefore, we hypothesized that the PP-InsPs could be interfering with the transcriptomic response to nutrient limitation. In this context, we conducted an RNA-seq analysis of wild-type and *vip1-1* cultures under control and nutrient limitation. In *C. reinhardtii*, the N assimilation involves a complex regulatory network comprising ammonium and nitrate sensors which induce transcription of the required genes such as membrane transporters and reduction enzymes (Calatrava et al., 2023). Nitrate is incorporated to the cells through the NRT2 transporters that require the accessory protein NAR2, to be later reduced by the nitrate reductase NIT1/NIA1 and nitrite reductase NII1 to produce ammonium, while ammonium uptake is produced through AMT1 transporters. We used ammonium in our culture medium as the N source, which resulted in no significant differences in the expression of nitrate assimilation related genes between strains at control condition (Figure 4A). However, when nutrient limiting conditions were reached, we found that genes necessary for nitrate uptake and nitrate reduction lacked their activation response in the *vip1-1* cells as they do in WT under the same conditions (Figure 4A). In fact, *vip1-1* slows its growth under nitrate as a source of N compared to WT (Supplementary figure 2). Even though we used general nutrient limitation conditions, *vip1-1* cells seem to display impairments in nitrogen depletion responses specifically. Current knowledge in nitrogen scavenging response in *C. reinhardtii* point to an upregulation in several genes (Park et al., 2015; Schmollinger et al., 2014). We found no significant differences between WT and *vip1-1* in the mRNA levels of different genes related to this process: ammonium transporters *amt1.2*, *amt1.5* and *amt1.8*, bispecific nitrate/nitrite transporters *nar1.2*, *nar1.6*, *nrt1.1*, and *nrt2.6*, glutamine synthetases *gln1*, *gln2*, *gln3* and gln4 and glutamine oxoglutarate aminotransferases *gsn* and *gsf* (Supplementary table II). In fact, glutamine synthetase (GS) activity was also measured under control conditions and nutritional deficiency in both strains (Supplementary figure 3). Under control conditions, GS activity was very similar in both strains; however, after prolonged cultivation GS activity was significantly higher in *vip1-1* compared to WT. This result points to an aberrant regulation of nitrogen scavenging that we further investigated. In this sense, we found genes lacking their activation response in nutrient deficiency in the *vip1-1* strain (*nrt2.1*, *nrt2.2*, *nar2*, *nit1* and *nii1*) (Figure 4A). These are all genes regulated by the transcription factor NIT2. Although none post-translational regulation has been reported to *nit2*, it is already known that the zinc finger protein NZF1 controls its expression modifying the 3’UTR length of the *nit2* transcript (Higuera et al., 2014; Sakuraba et al., 2022). We did not find significant differences in the mRNA accumulation levels of *nzf1 or nit2* genes (Supplementary table II), which suggests that PP-InsPs might be either acting together with the NZF1 protein controlling *nit2* at the translational level, or after *nit2* translation, mediating NIT2 activity as a transcription factor. In order to validate these findings, the activity of the nitrate reductase was assayed in the wild-type and *vip1-1* strains under control and nutrient deficiency conditions (Figure 4B). The results showed a significant reduction in the activity of this enzyme in the nutrient-limited *vip1-1* culture compared to the wild-type, which is consistent with the lack of mRNA accumulation of the nitrate reductase gene in *vip1-1* under these conditions (Figure 4B). Together, these results point to an important role of the PP-InsPs in nitrogen and possibly other nutrients limitation at the transcriptional level, facilitating the NIT2-driven transcriptomic response.

**Figure 4:**
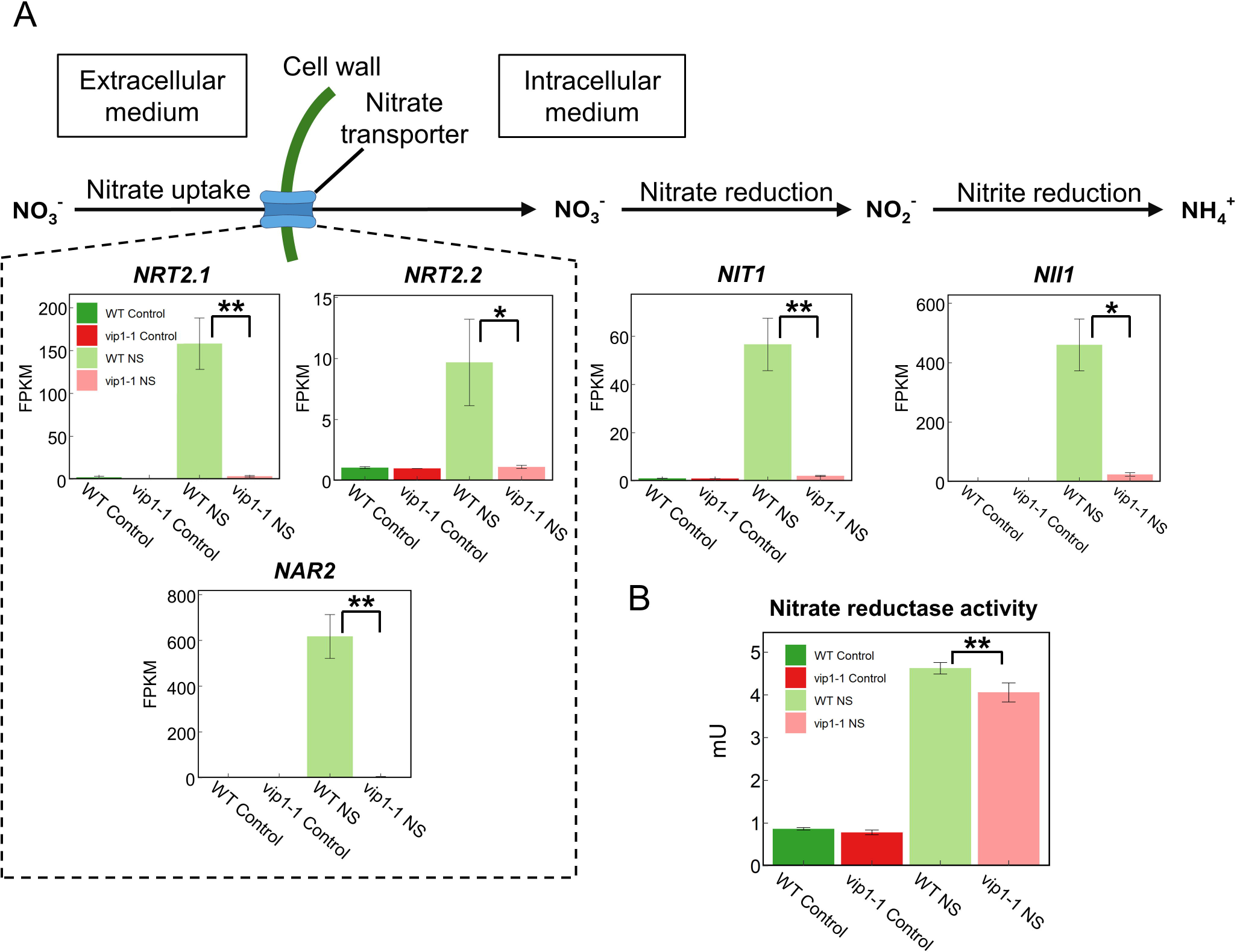
Diagram of nitrate assimilatory pathway (A), including gene expression of related genes. *NRT2.1* (Cre09.g410850) and *NRT2.2* (Cre09.g410800) encode nitrate transporters, both requiring the accesory protein encoded by *NAR2* (Cre09.g410900). *NIT1* (Cre09.g410950) encodes a nitrate reductase and *N/11* (Cre09.g410750) a nitrite reductase. Three independent biological replicates were analyzed in each condition (C; conrol and NS; nutritional stress) except from WT NS, where two independent biological replicates were analyzed. Error bars indicate mean±SE. B, nitrate reductase activity assay in each culture condition. Error bars indicate mean±SD. Asterisks indicates significant differences(* indicates p < 0.05, ** indicates p < 0.01).

## Discussion

Prolonged cultivation causes nutrient limitation and stress in microalgal cultures but these conditions sometimes are part of the scaling up process which is fundamental for their biotechnological use. In fact, there are some microalgal strains that need to be grown on specific PBRs with medium uptake that have restrictive volumes and increase the cost of obtaining microalgal biomass (Wan Mahari et al., 2022). For this reason, it is important to understand which controlling mechanisms underlie the resilience of microalgal cultures under limited nutrient conditions. This study intended to understand the importance of PP-InsPs in nutrient limitation responses in the model single-cell microalga *Chlamydomonas reinhardtii*, using *vip1-1* mutant (Couso et al., 2016).

In this context, we found that the inositol pyrophosphates-deficient strain *vip1-1* displayed a very interesting phenotype with higher chlorophyll content than the wild-type strain under nutritional deficiency (Figure 1A and B). However, chlorophyll fluorescence measurements found less photosynthetic activity and an impaired recovery capacity in the mutant compared to WT cells, specially under nutritional deficiency after 21 days grown in batch (Figure 1C and D). This phenotype has been shown by stay-green mutants in plants, in which mutations in genes involved in chlorophyll degradation were identified, leading to higher chlorophyll content and reduced photosynthetic capacity (Balazadeh, 2014). However, we did not find transcriptomic evidence regarding PP-InsPs affecting chlorophyll synthesis or breakdown, which hints that chlorophyll homeostasis must be affected by PP-InsPs in a post-transcriptional manner. In fact, we previously found an important deregulation of the phosphorylation patterns of many proteins belonging to PSII apparatus, including chlorophyll binding proteins such as CP43 and CP47 and several proteins belonging to the light harvesting antenna (Couso et al., 2021). Chlorophyll binding proteins have been reported to have an important interconnection in the degradation of chlorophylls in leaf senescence (Hörtensteiner and Feller, 2002). This chlorophyll degradation is induced by senescence in plants, but by nitrogen deprivation in microalgae like *C. reinhardtii*, although its regulation is yet to be elucidated (Chen et al., 2019; Willows et al., 2023). All these data indicate that PP-InsPs signaling is involved in the chlorophylls turnover by affecting the stability of the PSII. Also, the reduction in photosynthetic activity might be related to this chlorophyll accumulation affecting structure and/or function of the photosynthetic apparatus, or also to impairments in nitrogen scavenging mechanisms in *vip1-1*.

The chlorophyll binding proteins account for 20% of the cellular nitrogen, and these proteins can only be recycled once chlorophylls have been released and degraded in plants (Wang and Grimm, 2021). Therefore, the observed chlorophyll accumulation in *vip1-1* (Figure 1A, 1B) suggests that inositol pyrophosphates are required for the nitrogen scavenging mechanisms related to chlorophyll and chlorophyll binding proteins recycling.

In fact, the metabolic profiling revealed that amino acids involved in the arginine biosynthetic pathway were reduced in the absence of inositol pyrophosphates only in nutrient deficiency conditions (Figure 3B). Arginine levels have been reported to participate in the signaling network responsible for post-transcriptional control in N-deprived *C. reinhardtii* cultures (Monteiro et al., 2023). Interestingly, the decrease showed by *vip1-1* in most amino acids during nutrient limitation, especially the arginine biosynthesis-related ones (arginine, asparagine, aspartate and glutamate), supports the reduced chlorophyll and chlorophyll binding proteins recycling discussed above.

Electrodense bodies inside vacuoles displayed by TEM images in the *vip1-1* cells were observed since the beginning of the experiment but were especially visible after 14 days (Figure 2E). This might be a consequence of malfunctioning recycling process, which also correlates with lower amino acid accumulation under nutritional deficiency following misregulated intracellular nitrogen scavenging. In Couso et al., 2021, we reported aberrant ATG8 levels in *vip1-1* after rapamycin treatment that points to a deregulation of autophagy in the mutant under nutritional stress conditions when TOR activity is highly reduced. Here, we also observed an impaired transcriptomic response involving transcriptional activation of key genes in nitrate scavenge, e.g., nitrate transporters *nrt2.1*, *nrt2.2*, *nar2*, nitrate reductase *nit1* and nitrite reductase *nii1*, which was confirmed with a nitrate reductase assay as an alternative analytical method (Figure 4A, 4B, Supplementary table II). The transcriptomic response to nitrogen deprivation in *C. reinhardtii* involves several ammonium and nitrate transporters, as well as nitrate reductase, nitrite reductase and GS/GOGAT cycle enzymes (Park et al., 2015; Schmollinger et al., 2014). Nevertheless, only genes regulated by the transcription factor NIT2 were not activated under nutritional deficiency in the mutant, even though we did not find significant differences in *nit2* mRNA accumulation. The regulation of *nit2* is not well understood yet although NZF1 activity has been reported to affect its mRNA 3’UTR length (Higuera et al., 2014). Remarkably, this type of gene expression regulation has been found to be exerted by PP-InsPs in yeasts (Sanchez et al., 2019). These findings propose that these molecules must participate in nitrogen limitation responses controlling NIT2 activity, possibly modulating its 3’UTR length, and signaling for intracellular nutrient recycling. In this context, the low availability of free amino acid levels observed in *vip1-1* under nutrient stress (Figure 3A, 3B, Supplementary table1) lines up with the malfunctions in nitrogen scavenging and recycling suggested by our data.

Interestingly, TEM images showed that *vip1-1* cells were surrounded by a double plasmatic membrane (Figure 2F) after 21 days in batch cultivation. In *C. reinhardtii*, cells sense nutritional availability to either proceed with cell division or enter a quiescence state (Cross and Umen, 2015; Takeuchi and Benning, 2019). This is evaluated during G1 phase, when a cell is committed to perform at least one S/M cycle if it grows past a certain size but arrests its cell cycle if growth has not surpassed this “commitment” point due to nutrient scarcity or other environmental factors (Cross and Umen, 2015). After mitosis, daughter cells remain inside the cell wall of the mother cell, and secretion of lytic enzymes is required to perform hatching (Kubo et al., 2009). The regulatory network where PP-InsPs take part might be interfering with the adequate cell cycle progression in *C. reinhardtii*, which has been previously described for S phase in yeasts (Banfic et al., 2013). In fact, the transcription factor NIT2 shares the RWP-RK domain with MID, which determines the mating type of *C. reinhardtii* cells and responds to N deficiency (Innami et al., 2022; Lin and Goodenough, 2007; Sakuraba et al., 2022). Therefore, the PP-InsPs might be necessary for the proper activity of MID in the same way that they are for NIT2, as previously discussed. Given these findings, the PP-InsPs could be integrating nitrogen deprivation responses and cell cycle control with nutrient availability, potentially through transcription regulation and possibly post-transcriptional regulation of transcription factors.

## Acknowledgements

This research was supported by the Ministerio de Ciencia e Innovación TED2021-129409A-I00 and PID2022-136633OA-I00 grants awarded to IC. RBG was also awarded as a FPU22/00688 fellow by Ministerio de Educación.

## Author contributions

RBG, MEGG, and IC contributed to planning and experimental design; RBG, MEGG, JMP and IC performed experiments and data analysis; and RBG, MEGG, JMP and IC wrote the manuscript.

## Data availability statement

The raw and processed RNA-seq data that support the findings of this study are openly available in the Gene Expression Omnibus (https://www.ncbi.nlm.nih.gov/geo/) repository and can be accessed with the Identifier GSE276249. The raw RNA-seq data are openly available at the Sequence Read Archive (https://www.ncbi.nlm.nih.gov/sra) and can be accessed with the identifier PRJNA1156089. The raw and processed metabolomic data can be found in the MetaboLights repository (https://www.ebi.ac.uk/metabolights/) using the identifier MTBLS10779.

## Supporting information

Supplementary Figure 1: Starch levels in WT and *vip1-1* under control conditions, 14 days and 21 days in batch culture. Error bars indicate mean±SD, asterisks indicate significant differences evaluated using the Wilcoxon signed-rank test (* indicates p < 0.05).

Supplementary Figure 2: Growth curves of WT and *vip1-1* using NO_3_^-^ as N source. Error bars indicate mean±SD.

Supplementary figure 3: Glutamine synthetase activity assay in WT and *vip1-1* under control conditions and 21 days in batch culture. Error bars indicate mean±SE, asterisks indicate significant differences (* indicates p < 0.05, ** indicates p < 0.01).

Supplementary table I: Metabolite levels of wild-type and *vip1-1* strains. Values are expressed as log2FoldChange in control conditions (24 hours) and nutritional stress, NS (14 days). P-value was computed using the Wilcoxon signed-rank test and corrected with the Benjamini-Hochberg method. Significative differences are highlighted using light red or blue for the p-value cell if the increase or decrease in the mutant strain is less than 2-fold (log2FC less than 1 or -1), respectively. Dark red or blue was used if the increase or decrease in the mutant strain is bigger than 2-fold (log2FC bigger than 1 or -1), respectively.

Supplementary table II: RNA-seq analysis of nitrogen assimilation related genes. Colored blue log2(Fold Change) represents significative downregulated genes in *vip1-1* (adjusted p-value < 0.05) compared to WT while colored red log2(Fold Change) represents significative upregulated genes in *vip1-1* (adjusted p-value < 0.05) compared to WT.

## Notes

https://www.ncbi.nlm.nih.gov/geo/query/acc.cgi?acc=GSE276249

https://www.ebi.ac.uk/metabolights/editor/MTBLS10779/descriptors

### Competing Interest Statement

The authors have declared no competing interest.

