## Supplementary figures and images for "Inositol polyphosphates regulate resilient mechanisms in the green alga *Chlamydomonas reinhardtii* to adapt to extreme nutrient conditions"

### Supplemental Figure 1

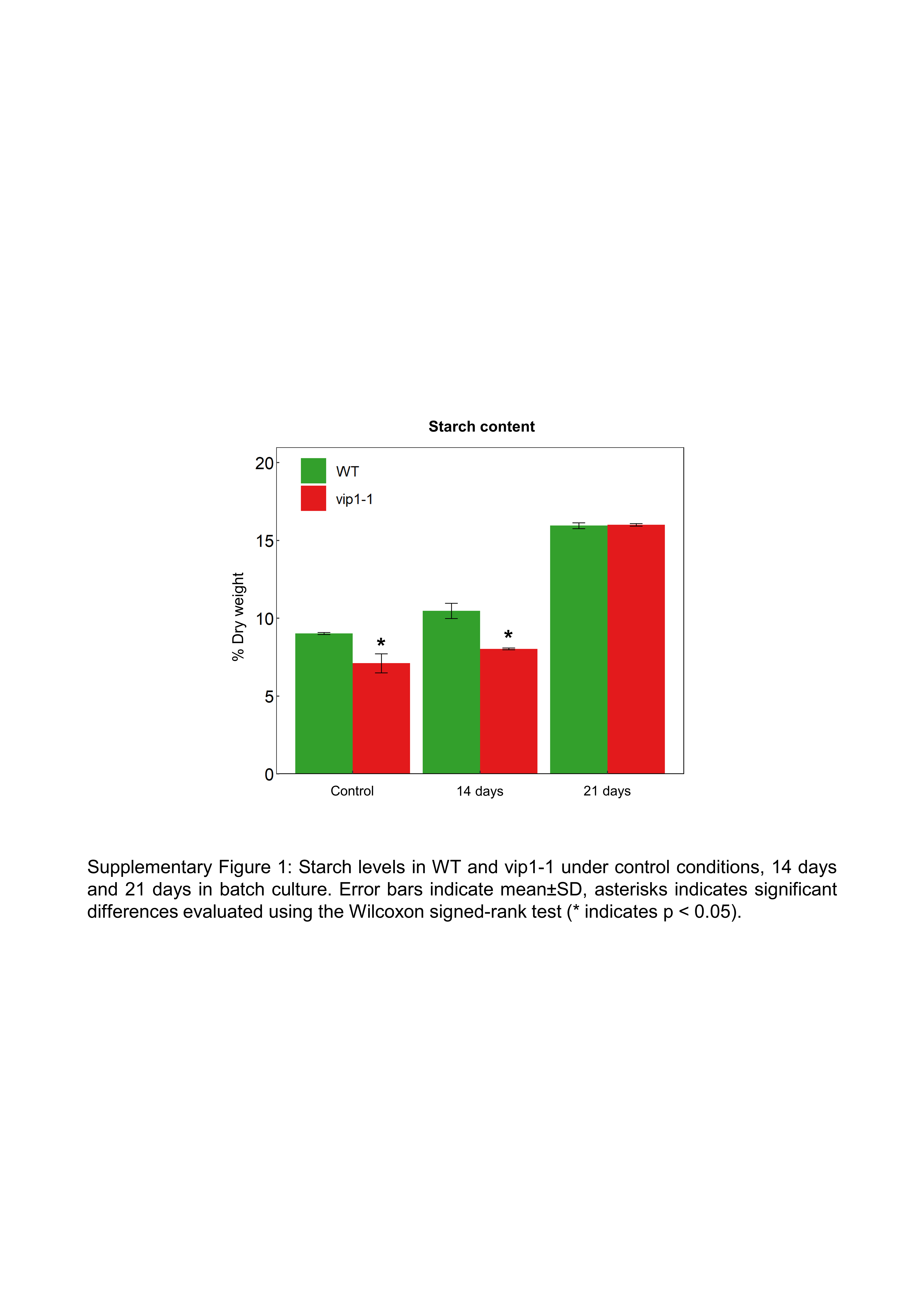

### Supplemental Figure 2

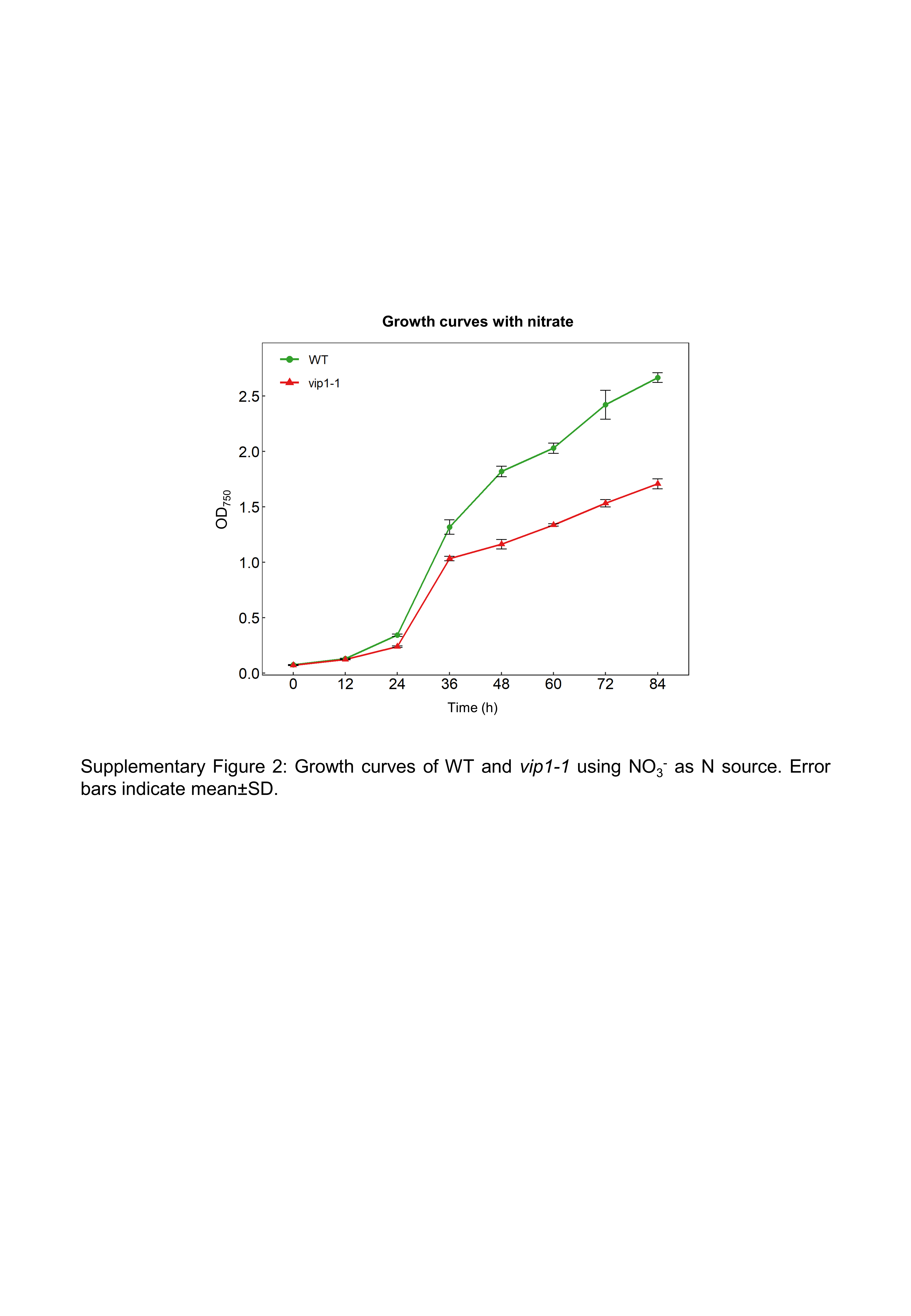

### Supplemental Figure 3

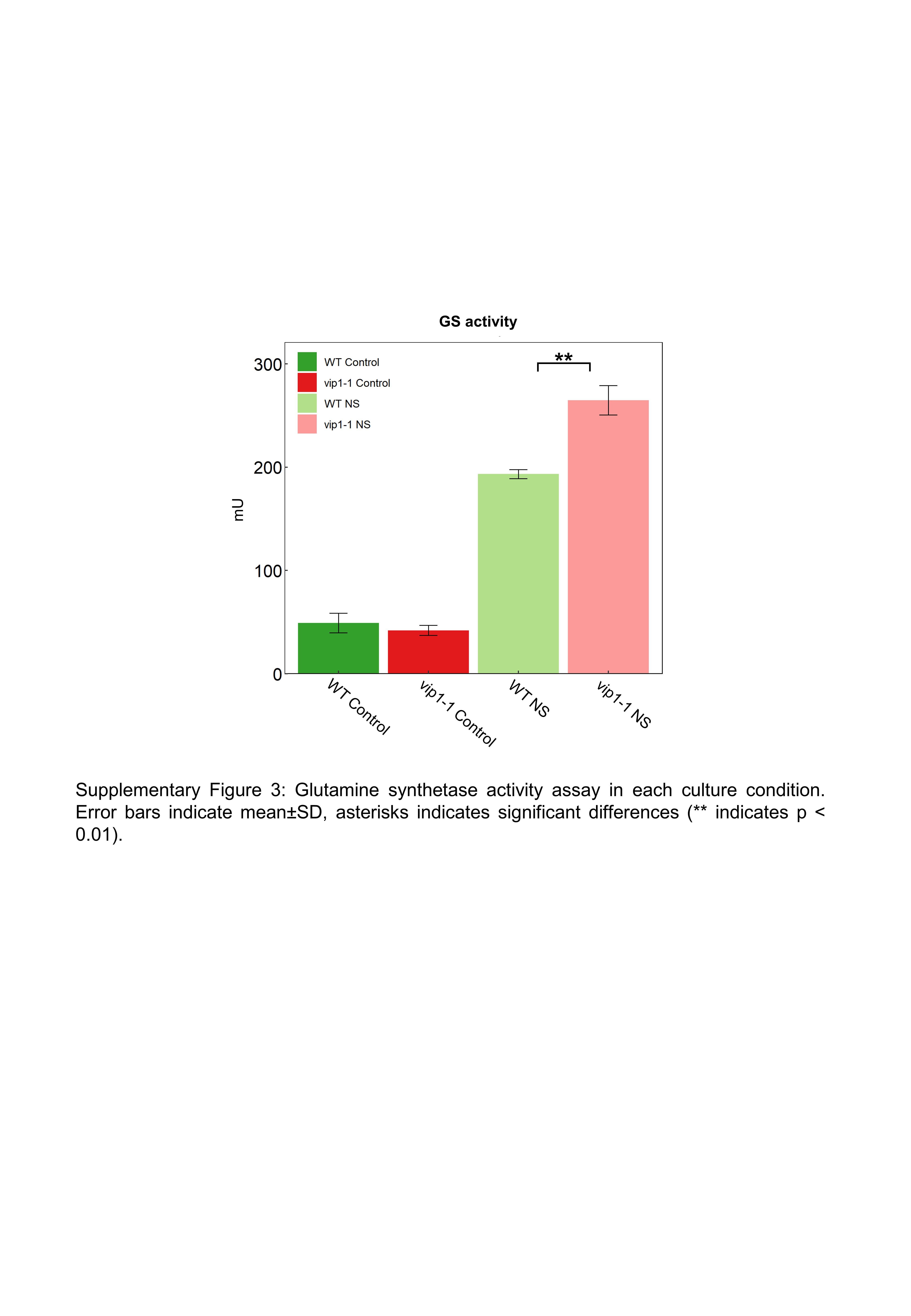
